# Complexity-Aware Simple Modeling

**DOI:** 10.1101/248419

**Authors:** Mariana Gómez-Schiavon, Hana El-Samad

**Author notes:** Email addresses (Mariana Gómez-Schiavon), (Hana El-Samad).

## Abstract

Mathematical models continue to be essential for deepening our understanding of biology. On one extreme, simple or small-scale models help delineate general biological principles. However, the parsimony of detail in these models as well as their assumption of modularity and insulation make them inaccurate for describing quantitative features. On the other extreme, large-scale and detailed models can quantitatively recapitulate a phenotype of interest, but have to rely on many unknown parameters, making them often difficult to parse mechanistically and to use for extracting general principles. We discuss some examples of a new approach — complexity-aware simple modeling — that can bridge the gap between the small‐ and large-scale approaches.

**Highlights:**

- Simple or small-scale models allow deduction of fundamental principles of biological systems
- Detailed or large-scale models can be quantitatively accurate but difficult to analyze
- Complexity-aware simple models can extract principles that are robust to the presence of unknown complex interactions

Graphical abstract

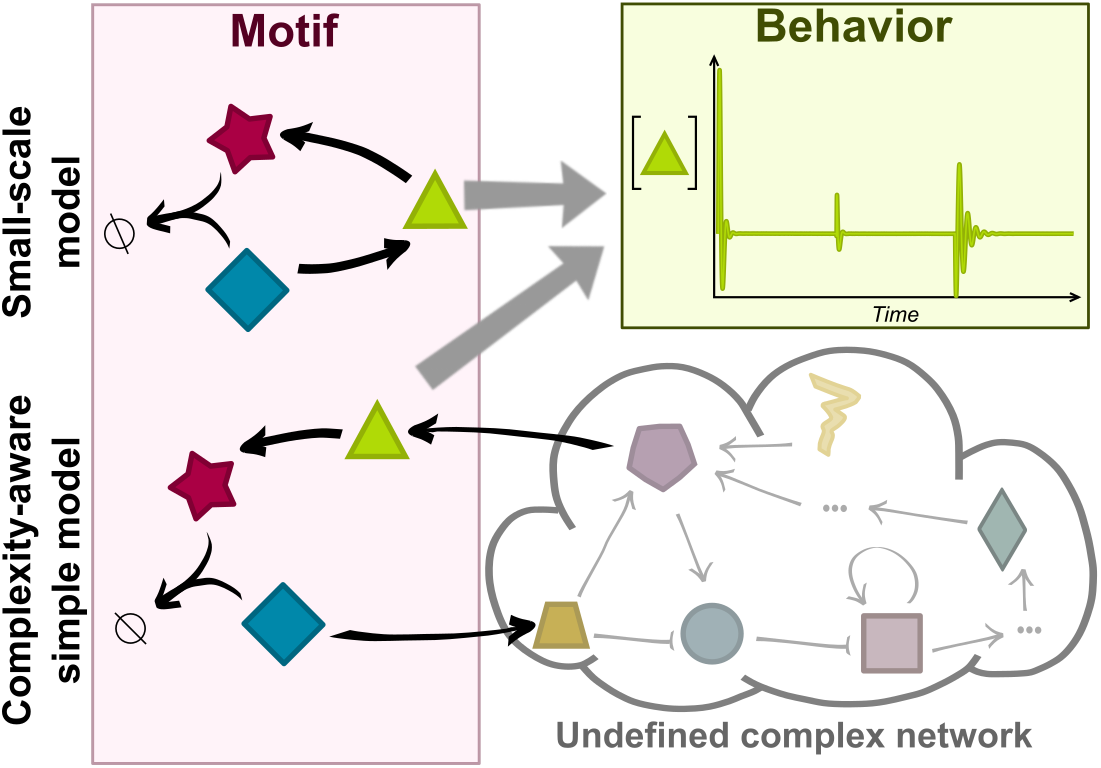

## Introduction

Mathematical models have long been the crutch for our intuition in many fields of science. Models have also rapidly become accepted tools in the biological sciences, used to organize data and knowledge, understand how biological phenomena arise from the collective action of components [1] and predict emergent organizational properties [2, 3].

In general, the most useful model for a particular process would depend on the specific question at hand as well as the information available (previous knowledge and attainable experimental data) [4, 5, 6]. As a result, a single model is rarely appropriate for all possible instances of a problem [7]. This said, modelers of biology have long argued (and continue to do so) about the most useful approach, creating some tension between the supporters of large‐ and small-scale models [8]. Detailed or large-scale models attempt to incorporate most or all the available information about a system that is being modeled, resulting in many components and interactions explicitly stated in the resulting model. These models are criticized for being poorly parametrized and not easily amenable to abstraction and general insight. On the other hand, simple or small-scale models actively seek to discern the minimal essential components and interactions required to explain a particular behavior. As a result, the quantitative predictive power of small-scale models is often questioned. All models, by definition, fail to incorporate every mechanistic detail of a biological system [9]. Simply stated, since models are necessarily approximations of reality, even the most elaborate model contains a set of assumptions, and any conclusion derived from this model would be dependent on the validity of these assumptions.

While keeping this in mind, we discuss some examples of “small-scale” models and “large-scale” models. We then introduce the potential hybridization of these two through an approach we call “complexity-aware simple modeling”. We discuss two recent examples of this promising approach.

### *Small-scale* models: The power of simplicity

The motivation behind small-scale models is that the most parsimonious set of components and their interactions that can explain a phenotype also provide the most power for unraveling its underlying requirements. These models provide a major benefit: by using a small number of components, the number of unknown parameters is minimal and their associated assumptions are tractable. This greatly facilitates interpretation and provides an opportunity for vetting the generality of conclusions. As a result, small-scale models are often associated with the quest for uncovering principles.

In support of this notion, many concepts that are deeply embedded in our current knowledge of biological circuits result from small-scale models (see [10]). These include prominent principles such as the need for positive feedback for multistability [11], and negative feedback and time delays to produce oscillations [12]. Such simple but powerful guiding principles have been crucial for the study of many biological systems, ranging from circadian rhythms [13, 14, 15] to cell cycle regulation [16, 17, 18], and have led to profound insights into these complex systems.

While many small-scale models are derived with a biological system in mind, some are constructed to probe the general requirements of a broad biological property. For example, small-scale models have been used to pinpoint specific structural attributes of the biochemical networks that produce “absolute concentration robustness” (ACR). An ACR network is one in which one of the molecular species maintains a constant concentration, irrespective of fluctuations of other circuit components. A simple model of two molecular species *A* and *B*, present at a total concentrate *A* + *B* = *θ*, interconverting along the two simple reactions *A* + *B* → 2*B* and *B* → *A* (at rates *α* and *β* respectively) shows ACR in that the concentration of *A* is constant (at *β*/*α*) irrespective of total concentration *θ*. This example motivated the development of a broad theory for defining large classes of ACR networks [19, 20] that can produce this property irrespective of biochemical parameter values.

Often however, the most meaningful understanding derives from a convergence of the general investigations of “principles” with the focused investigations of a concrete biological network. Unraveling how frog oocytes implement an irreversible differentiation switch was accomplished through a keen interest in this biological question as well as exploration of the properties of ultrasensitivity and positive feedback using simple models [21, 22, 23]. Another example is that of perfect adaptation in which a functional quantity of a biological circuit can maintain a steady-state value that is constant despite a perturbing input. A multi-decade interest in understanding how bacteria implement perfect adaptation in chemotaxis [24, 25, 26, 27, 28, 29] led to a compelling formulation of this problem, with renewed interest generated by the identification of perfect adaptation in other systems [30, 31, 32]. Here again, insights gained from simple models of biological systems that feature perfect adaptation (see [33]) converged with general inquiries about motifs and topological features that can generate such a property (see [34]) to produce a meaningful and deep understanding. In particular, many of these studies converged on the use of integral feedback control, which was mathematically demonstrated decades ago to ensure perfect adaptation in the field of control theory [26, 35, 36].

Despite the success of small-scale and simple models, biology itself is neither small-scale nor simple. And, when using small-scale models to describe local nodes of a bigger biological network or to simplify elaborate interactions, we should continuously challenge our conclusions by asking about the effects of the surrounding complexity. What happens if the network motif that ensures perfect adaptation has an extra link or is connected to another network, or if the positive feedback that implements a switch is also entangled in a negative feedback loop? Each of these cases would have to be explored thoroughly, building our understanding from the bottom-up.

### *Large-scale* models: Embracing biological complexity

The premise of large-scale models is that all components and interactions that comprise a system might be needed in order to reproduce its quantitative behavior accurately. The construction and simulation of such models that are faithful to details and complexity is now facilitated by acceleration in experimental data collection and growth in computational power (e.g. see [37]).

At the extreme of this spectrum are studies that attempt whole-cell modeling, seeking to describe how the phenotype arises from the genotype by accounting for all genes/proteins and interactions in a cell (i.e. human pathogen *Mycoplasma genitalium*), integrating multiple sources of data, as the transcriptome, proteome, and metabolome in a condition of interest, as well as more general properties of the cell, such as mass, geometry, and cell-cycle state [38, 39]. The resulting models have so far included hundreds of variables and thousands of parameters whose values have to be mostly assumed. Insights generated by these models include the identification of new gene functions and the prediction of biological processes not directly accessible by existing experimental measurements [38].

Large-scale models also arise from efforts to reconstruct cellular networks in an unbiased way (top-down) from high-throughput data [40]. These reconstructions have proven to be useful to provide an overview of cellular connectivity, but the analyses of the resulting models have often focused on isolating a few structural components and interactions associated with a phenotype of interest [41].

A third general approach for large-scale modeling is one in which the complexity is built stepwise, first by building simple models and then embedding them in a more elaborate physiological reality. For example, Spiesser *et al*. [42] presented a multiscale simulation platform to integrate an osmostress response model with its physiological context (e.g. cell division cycle), as well as the cell-to-cell variation expected in a cellular culture. This model revealed previously under-estimated features that are dependent on population dynamics, such as partial synchronization during osmoadaptation [42].

Overall, while large-scale models are undeniably closer to biological reality, the task of interpreting their findings remains difficult. The many parameters of these models, most of which are poorly measured or not measured at all, makes it difficult to differentiate conclusions and predictions that are dependent on parameter choices from those that are robust and general.

## A new approach: Accounting for complexity without getting entangled in it

Recent years have seen the emergence of an exciting modeling approach, which we call here “complexity-aware simple modeling”. The goal is to preserve the small-scale modeling approach for representing a biological process, but without ignoring the complexity surrounding it. In fact, the point is to identify the largest and most complex class of interactions, which when connected to the simple model, fail to perturb its behavior. In this framework, the biological process of interest is modeled with the resolution needed but the surrounding complexity (e.g. connected networks) is deliberately kept undefined or defined by the most abstract representation possible. Statements about the behavior of the system of interest are then formulated and demonstrated to hold even in the presence of the unmodeled interactions (Figure 1).

**Figure 1:**
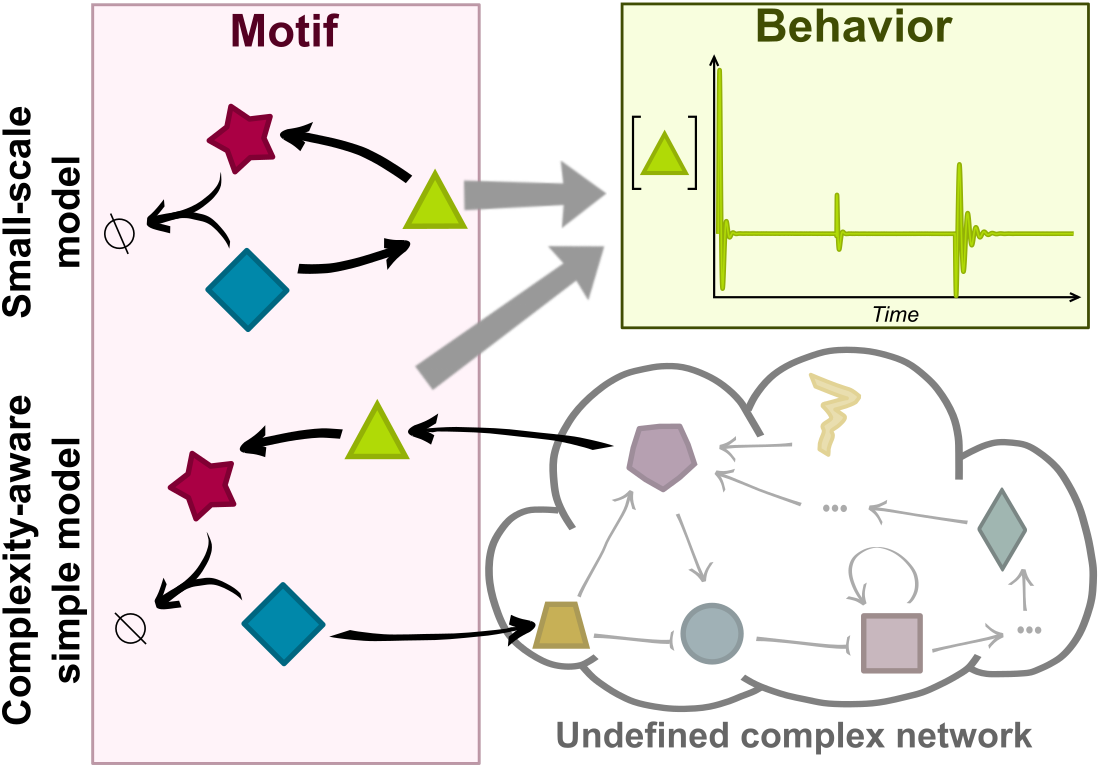
Complexity-aware simple models. consider a small number of components and interactions necessary to explain a behavior of interest. They also include an abstract representation of the largest class of complex interactions, which when connected to the simple model, have a minimal impact on its behavior.

A representative example of this approach asked whether there is a simple biochemical motif (with its corresponding simple model) that can achieve integral feedback when connected to an arbitrarily complex network (with any number of components and interactions, as well as undefined parameter values) [43]. The result was the so-called “antithetic motif”, where two molecular species bind to each other and annihilate each others function through this binding. Now, imagine that one of the “antithetic” molecular species controls the input of the complex network while the other is produced by the output of the same network. In this case, it can be mathematically demonstrated that the steady-state value of the network’s output perfectly adapts regardless of any step perturbation inside this network (Figure 1). The antithetic motif used in this configuration therefore implements integral feedback action. One requirement of this adaptation is that the only source of decay for the two molecules of the antithetic motif is their mutual annihilation, not their individual degradation or inactivation. While such perfect adaptation holds for an arbitrarily complex network connected to the antithetic motif, it is of course contingent on the network being responsive to the input from this motif. Still, this is a remarkable property with two main implications. First, it provides a recipe for building a simple “adduct” to a very complex network that makes it perfectly adapting. Second, when this antithetic motif is found in endogenous biological networks, we can isolate the motif, ignore the rest and declare the network perfectly adapting without the need to detail its complexity in order to infer the perfect adaptation property. A similar approach has been implemented to identify other integral control motifs [44] and to prescribe a general and robust cell fate reprogramming strategy [45].

A variation on this theme seeks to find bounds on behavior for classes of systems that share a small number of parts but can be arbitrarily different in others. For instance, being cognizant that all molecular interactions in cells are probabilistic, it is possible to define general relationships and bounds on the cell-to-cell variability in the antithetic motif (explained above) that hold irrespective of any complex network connected to it. Assuming stochastic birth, death, and binding reactions of the molecules in this motif, algebraic expressions based on relating average abundances and covariances for the two antithetic species can be combined with simple mathematical properties of normalized covariances to derive an appropriate bound on the fluctuations of these species. This approach showed a fundamental trade-off: molecular fluctuations in the counts of free molecules have to increase if a higher efficiency of binding between the antithetic molecules (and formation of their bimolecular complex) is desired [46]. This bound holds irrespective of, and cannot be alleviated by, connectivity to any network of arbitrary size or complexity. Therefore, this relationship is only based on a few specified interactions and is invariant to any networks in which those interactions might be embedded. This idea can further be exploited to rule out the plausibility of specific classes of interactions for an underlying biological process given experimental data. For example, if one is hypothesizing the presence of an antithetic motif, but experimental measurements of fluctuations and complex formation efficiency are outside the general bound delineated by their theoretically determined relationship, one can efficiently rule out the involvement of this motif irrespective of other un-characterized components present in the network. Such an approach was productively used in a different context to determine that a common class of gene expression models in which protein synthesis is proportional to mRNA levels cannot account for experimental single cell measurements in *E. coli* [47, 48]. This determination was based on the fact that the measured values of covariance metrics between mRNA and protein differ significantly from predicted relationships. Here again, predicted values are based on a simple model but are invariant to the potential complex connectivity of this model.

## Final thought

Systems biology is often thought of as “the tool to unravel black boxes” [49]. We pose here the question of whether, when modeling biological systems, it is sometimes more productive to deliberately keep some boxes closed through what we have called “complexity-aware simple models”, or other approaches that adopt a similar philosophy. We find this idea appealing, and advocate for considering its implications. Might it be a fruitful way to approach the coarse-graining that is needed to traverse the different scales of biological organization? Might it a useful replacement for detailed descriptions of certain processes in whole cell models?

At the same time, we caution that only a few examples of the success of this approach exist and that there is still no clear disciplined way to implement such analyses in a general sense. We also caution that models are used for a variety of reasons and for asking different questions [5]. Therefore, we must continue to unabatedly define models at a resolution that enables them to be useful tools for answering these questions. Fundamentally, we aim our discussion of complexity-aware simple models to provide some food for thought and hopefully a subject for a vigorous scientific debate.

## Acknowledgements

This work was supported by the Paul G. Allen Family Foundation and the National Science Foundation (NSF-MCB 1715108) to H.E.S. Hana El-Samad is a Chan Zuckerberg Biohub investigator.

## References

[1] Wolkenhauer O. Why model? Frontiers in Physiology 2014;5(January):1–5. doi:10.3389/fphys.2014.00021. • A short review of modeling approaches for biological systems ranging from small-scale to large-scale models, with their advantages and limitations.

[2] Gunawardena J. Biology is more theoretical than physics. Molecular Biology of the Cell 2013;24(12):1827–9. doi:10.1091/mbc.E12-03-0227.

[3] Tsigkinopoulou A, Baker SM, Breitling R. Respectful Modeling: Addressing Uncertainty in Dynamic System Models for Molecular Biology. Trends in Biotechnology 2017;35(6):518–29. doi:10.1016/j.tibtech.2016.12.008.

[4] Kirk P, Thorne T, Stumpf MP. Model selection in systems and synthetic biology. Current Opinion in Biotechnology 2013;24(4):767–74. doi:10.1016/j.copbio.2013.03.012.

[5] Gunawardena J. Models in biology: accurate descriptions of our pathetic thinking’. BMC Biology 2014;12(1):29. doi:10.1186/1741-7007-12-29. • An engaging discussion of the concept of mathematical models in biology, with particular emphasis on the role of the underlying assumptions. The author analyzes several modeling examples and suggests general guidelines to build biological models.

[6] Silk D, Kirk PDW, Barnes CP, Toni T, Stumpf MPH. Model Selection in Systems Biology Depends on Experimental Design. PLoS Computational Biology 2014;10(6):e1003650. doi:10.1371/journal.pcbi.1003650.

[7] Evans MR, Grimm V, Johst K, Knuuttila T, de Langhe R, Lessells CM, et al. Do simple models lead to generality in ecology? Trends in ecology & evolution 2013;28(10):578–83. doi:10.1016/j.tree.2013.05.022.

[8] Mellis IA, Raj A. Half dozen of one, six billion of the other: What can small‐ and large-scale molecular systems biology learn from one another? Genome Research 2015;25(10):1466–72. doi:10.1101/gr.190579.115.

[9] Reeves GT, Fraser SE. Biological Systems from an Engineer’s Point of View. PLoS Biology 2009;7(1):e1000021. doi:10.1371/journal.pbio.1000021.

[10] Alon U. Network motifs: theory and experimental approaches. Nature reviews Genetics 2007;8(6):450–61. doi:10.1038/nrg2102.

[11] Thomas R, Kaufman M. Multistationarity, the basis of cell differentiation and memory. I. Structural conditions of multistationarity and other nontrivial behavior. Chaos: An Interdisciplinary Journal of Nonlinear Science 2001;11(1):170. doi:10.1063/1.1350439.

[12] Novak B, Tyson JJ. Design principles of biochemical oscillators. Nature Reviews Molecular Cell Biology 2008;9(12):981–91. doi:10.1038/ nrm2530.

[13] Sehgal A, Rothenfluh-Hilfiker A, Hunter-Ensor M, Chen Y, Myers MP, Young MW. Rhythmic Expression of timeless: A Basis for Promoting Circadian Cycles in period Gene Autoregulation. Science 1995;270(5237):808–10. doi:10.1126/science.270.5237.808.

[14] Tomita J, Nakajima M, Kondo T, Iwasaki H. No Transcription-Translation Feedback in Circadian Rhythm of KaiC Phosphorylation. Science 2004;doi:10.1126/science.1102540.

[15] Nohales MA, Kay SA. Molecular mechanisms at the core of the plant circadian oscillator. Nature Structural & Molecular Biology 2016;23(12):1061–9. doi:10.1038/nsmb.3327.

[16] Tyson JJ, Chen KC, Novak B. Sniffers, buzzers, toggles and blinkers: dynamics of regulatory and signaling pathways in the cell. Current Opinion in Cell Biology 2003;15(2):221–31. doi:10.1016/S0955-0674(03)00017-6.

[17] Verdugo A, Vinod PK, Tyson JJ, Novak B. Molecular mechanisms creating bistable switches at cell cycle transitions. Open Biology 2013;3(3):120179–. doi:10.1098/rsob.120179.

[18] Ferrell JE. Feedback loops and reciprocal regulation: recurring motifs in the systems biology of the cell cycle. Current opinion in cell biology 2013;25(6):676–86. doi:10.1016/j.ceb.2013.07.007.

[19] Shinar G, Feinberg M. Structural Sources of Robustness in Biochemical Reaction Networks. Science 2010;327(5971):1389–91. doi:10.1126/science.1183372.

[20] Enciso GA. Transient absolute robustness in stochastic biochemical networks. Journal of The Royal Society Interface 2016;13(121):20160475. doi:10.1098/rsif.2016.0475. • This papers studies the dynamical behavior of systems exhibiting Absolute Concentration Robustness in the presence of biochemical noise to probe when and how noise disrupts the deterministic system.

[21] Ferrell JE, Ha SH. Ultrasensitivity part I: Michaelian responses and zero-order ultrasensitivity. Trends in Biochemical Sciences 2014;39(10):496–503. doi:10.1016/j.tibs.2014.08.003. • This and the following reviews in the series [22, 23] explore our understanding of ultrasensitivity in dynamical systems, mostly using small-scale models.

[22] Ferrell JE, Jr, Ha SH. Ultrasensitivity part II: multisite phosphorylation, stoichiometric inhibitors, and positive feedback. Trends in Biochemical Sciences 2014;39(11):556–69. doi:10.1016/j.tibs.2014.09.003.

[23] Ferrell JE, Ha SH. Ultrasensitivity part III: cascades, bistable switches, and oscillators. Trends in Biochemical Sciences 2014;39(12):612–8. doi:10.1016/j.tibs.2014.10.002.

[24] Berg HC, Brown DA. Chemotaxis in Escherichia coli analysed by Three-dimensional Tracking. Nature 1972;239(5374):500–4.

[25] Barkai N, Leibler S. Robustness in simple biochemical networks. Nature 1997;387(6636):913–7. doi:10.1038/43199.

[26] Yi TM, Huang Y, Simon MI, Doyle J. Robust perfect adaptation in bacterial chemotaxis through integral feedback control. Proceedings of the National Academy of Sciences 2000;97(9):4649–53. doi:10.1073/pnas.97.9.4649.

[27] Dufour YS, Fu X, Hernandez-Nunez L, Emonet T. Limits of Feedback Control in Bacterial Chemotaxis. PLoS’ Computational Biology 2014;10(6):e1003694. doi:10.1371/journal.pcbi.1003694.

[28] Frankel NW, Pontius W, Dufour YS, Long J, Hernandez-Nunez L, Emonet T. Adaptability of non-genetic diversity in bacterial chemotaxis. eLife 2014;3(October 2014):1–30. doi:10.7554/eLife.03526.

[29] Waite AJ, Frankel NW, Dufour YS, Johnston JF, Long J, Emonet T. Non-genetic diversity modulates population performance. Molecular Systems Biology 2016;12(12):895. doi:10.15252/msb.20167044.

[30] El-Samad H, Goff JP, Khammash M. Calcium homeostasis and parturient hypocalcemia: an integral feedback perspective. Journal of theoretical biology 2002;214(1):17–29. doi:10.1006/jtbi.2001.2422.

[31] Miller AJ, Smith SJ. Cytosolic nitrate ion homeostasis: Could it have a role in sensing nitrogen status? Annals of Botany 2008; 101(4):485–9. doi:10.1093/aob/mcm313.

[32] Muzzey D, Gómez-Uribe CA, Mettetal JT, van Oudenaarden A. A Systems-Level Analysis of Perfect Adaptation in Yeast Osmoregulation. Cell 2009;138(1):160–71. doi:10.1016/j.cell.2009.04.047.

[33] Somvanshi PR, Patel AK, Bhartiya S, Venkatesh KV. Implementation of integral feedback control in biological systems. Wiley In-terdisciplinary Reviews: Systems Biology and Medicine 2015;7(5):301–16. doi:10.1002/wsbm.1307.

[34] Ma W, Trusina A, El-Samad H, Lim WA, Tang C. Defining Network Topologies that Can Achieve Biochemical Adaptation. Cell 2009;138(4):760–73. doi:10.1016/j.cell.2009.06.013.

[35] Francis BA, Wonham WM. The internal model principle of control theory. Automatica 1976;12(5):457–65. doi:10.1016/0005-1098(76)90006-6.

[36] Bennett S. Development of the PID controller. IEEE Control Systems 1993;13(6):58–62. doi:10.1109/37.248006.

[37] Ji Z, Yan K, Li W, Hu H, Zhu X. Mathematical and Computational Modeling in Complex Biological Systems. BioMed Research Interna-tional 2017;2017:1–16. doi:10.1155/2017/5958321.

[38] Karr JR, Sanghvi JC, Macklin DN, Gutschow MV, Ja cobs JM, Bolival B, et al. A Whole-Cell Computational Model Predicts Phenotype from Genotype. Cell 2012;150(2):389–401. doi:10.1016/j.cell.2012.05.044. • An example of large-scale modeling.

[39] Karr JR, Takahashi K, Funahashi A. The principles of whole-cell modeling. Current Opinion in Microbiology 2015;27:18–24. doi:10.1016/j.mib.2015.06.004.

[40] Papin JA, Hunter T, Palsson BO, Subramaniam S. Reconstruction of cellular signalling networks and analysis of their properties. Nature Reviews Molecular Cell Biology 2005;6(2):99–111. doi:10.1038/nrm1570.

[41] Gat-Viks I, Shamir R. Refinement and expansion of signaling pathways: The osmotic response network in yeast. Genome Research 2007;17(3):358–67. doi:10.1101/gr.5750507.

[42] Spiesser T, Kühn C, Krantz M, Klipp E. The MYpop toolbox: Putting yeast stress responses in cellular context on single cell and population scales. Biotechnology Journal 2016;11(9):1158–68. doi:10.1002/biot.201500344. • A pipeline integrating multi-scale data to build large-scale quantitative models that account for cell-to-cell variability.

[43] Briat C, Gupta A, Khammash M. Antithetic Integral Feedback Ensures Robust Perfect Adaptation in Noisy Biomolecular Networks. Cell Systems 2016;2(1):15–26. doi:10.1016/j.cels.2016.01.004. arXiv:1410.6064. •• Prominent example of complexity-aware simple modeling, where an “antithetic” motif is demonstrated to provide perfect adaptation to an arbitrary network connected to it, even in the presence of biochemical noise.

[44] Briat C, Zechner C, Khammash M. Design of a Synthetic Integral Feedback Circuit: Dynamic Analysis and DNA Implementation. ACS Synthetic Biology 2016;5(10):1108–16. doi:10.1021/acssynbio.6b00014. •• This paper suggests a strategy for an “integral feedback” motif that achieves perfect adaptation and is robust to surrounding complexity.

[45] Del Vecchio D, Abdallah H, Qian Y, Collins JJ. A Blueprint for a Synthetic Genetic Feedback Controller to Reprogram Cell Fate. Cell Systems 2017;4(1):109–120.e11. doi:10.1016/j.cels.2016.12.001. •• This paper suggests a synthetic feedback control strategy to efficiently reprogram cell fate.

[46] Hilfinger A, Norman TM, Vinnicombe G, Paulsson J. Constraints on Fluctuations in Sparsely Characterized Biological Systems. Physical Review Letters 2016;116(5):1–5. doi:10.1103/PhysRevLett.116.058101. •• Example of complexity-aware simple modeling that identifies fundamental trade-offs in the stochastic behavior of a simple motif that holds irrespective of a complex connected network.

[47] Hilfinger A, Norman TM, Paulsson J. Exploiting Natural Fluctuations to Identify Kinetic Mechanisms in Sparsely Characterized Systems. Cell Systems 2016;2(4):251–9. doi:10.1016/j.cels.2016.04.002. •• An example where complexity-aware simple models are used to systematically test distinct mathematical models against experimental data and rule out classes of assumed biological interactions irrespective of surrounding complexity.

[48] Taniguchi Y, Choi PJ, Li GW, Chen H, Babu M, Hearn J, et al. Quantifying E. coli Proteome and Transcriptome with SingleMolecule Sensitivity in Single Cells. Science 2010;329(5991):533–8. doi:10.1126/science.1188308.

[49] Takors R, de Lorenzo V. Editorial overview: Microbial systems biology: systems biology prepares the ground for successful synthetic biology. Current Opinion in Microbiology 2016;33:viii–x. doi:10.1016/j.mib.2016.08.003.

